# Efficacy profiling of structurally diverse cannabinoid receptor ligands across transducer-coupling assays

**DOI:** 10.64898/2026.09.27.754774

**Authors:** Soo Jung Oh, Chrisotpher Lucaj, Kiera Truong, Ruiru Guo, Hayato Umaoka, Christos Iliopoulos-Tsoutsouvas, Maria Gerasi, Markos-Orestis Georgiadis, Lipin Ji, Spyros P Nikas, Alexandros Makriyannis, Hideaki Yano

## Abstract

The full efficacy range of ligands at CB1 and CB2 (i.e., inverse agonists, neutral agonists, partial agonists, full agonists, and high-efficacy agonists) has been reported collectively across numerous laboratories and assay platforms. Canonical Gi/o and β-arrestin signaling have each been suggested to shape physiological effects differently via different classes of cannabinoid ligands. These prior efficacy classifications, however, were obtained in multiple assay systems, often using indirect, amplified readouts and different reference agonists, making them difficult to compare across ligands. Here we profiled a structurally diverse panel of 22 cannabinoid ligands: synthetic cannabinoid receptor agonists (SCRAs), Δ⁹-tetrahydrocannabinol (Δ⁹-THC) analogues, endocannabinoid analogues, and antagonists/inverse agonists at CB1 and CB2 receptors using bioluminescence resonance energy transfer (BRET) assays, measuring Gαi1 engagement and β-arrestin 2 recruitment and normalizing every response to the reference full agonist CP55,940. At CB1, SCRAs and Δ⁹-THC analogues acted as high-efficacy agonists. At CB1, these high-efficacy agonists were also highly efficacious in β-arrestin 2 recruitment. Across the panel, the most consistent trend was a loss of agonist efficacy from CB1 to CB2. Among antagonists, inverse agonists suppressed constitutive CB1 activity more than neutral antagonists. Together, these CP55,940-normalized measurements provide a common efficacy scale and a framework for interpreting pharmacodynamics of CB1 and CB2 cannabinoid receptors.

## Introduction

Cannabis is among the most widely used psychoactive substances worldwide (World Drug Report, 2026) with a fast-growing rate of use (269% 2008 vs. 2022). Against this backdrop and with sustained illicit abuse of SCRAs (Caulkins, 2024), efficacy evaluation of diverse cannabinoid receptor ligands is becoming increasingly important, both for understanding abuse-related toxicity and for assessing therapeutic potential.

Cannabinoids exert wide-ranging central and peripheral effects that are mediated primarily through the endocannabinoid system (ECS), acting at two G protein-coupled receptors (GPCRs), CB1 and CB2. Both receptors canonically couple to inhibitory Gi/o proteins and, following agonist-induced phosphorylation, recruit β-arrestin, which drives receptor desensitization and internalization (Ibsen et al., 2017). Cannabinoids also act, to a lesser extent, through non-canonical targets including GPR55, TRPV1, and PPARs (Di Marzo and De Petrocellis, 2012).

CB1 is among the most abundantly expressed GPCRs in the central nervous system, where it mediates the psychoactive and both therapeutic- and abuse-relevant effects of cannabinoids (Zou and Kumar, 2018). CB2, by contrast, is predominantly peripheral and immune, mediating mainly immunomodulatory functions (Turcotte et al., 2016). This divergence is what makes subtype selectivity worth studying, with distinct therapeutic possibilities across agonists and antagonists at each receptor. CB2 selective agonists can be pursued for pain and inflammation without CB1 mediated psychoactivity, while CB1 agonists such as THC-mimetics dronabinol and nabilone for chemotherapy induced nausea and AIDS related anorexia, and the THC:CBD spray nabiximols for multiple sclerosis spasticity and neuropathic pain (Malan et al., 2003; Tsang and Giudice, 2016). On the antagonist side, there are peripherally restricted and neutral CB1 antagonists that preserve metabolic benefit: reducing adiposity, insulin resistance, and dyslipidemia, while sparing central receptors (Cinar et al., 2020). These same peripheral CB1 antagonists also show antifibrotic activity in preclinical models of liver and other organ fibrosis (Chen et al., 2024; Yang et al., 2024). CB2 antagonists and inverse agonists are the least clinically advanced but have shown preclinical anti-resorptive activity in models of bone loss and inflammatory bone disease, within CB2 bone biology in which agonists are also osteoprotective (Idris et al., 2005). Selectivity is difficult to achieve, however, because CB1 and CB2 share roughly 68% sequence identity across their transmembrane orthosteric-binding regions (Munro et al., 1993). As a result, many cannabinoids, including Δ⁹-tetrahydrocannabinol (Δ⁹-THC), bind both receptors with comparable affinity even as their localization and physiology diverge (Pertwee, 2008).

To this date, the cannabinoid research field has characterized CB1 and CB2 ligands spanning the entire efficacy range: inverse agonists, neutral antagonists, partial agonists, full agonists, and high efficacy agonists. The endocannabinoids anandamide and 2-arachidonoylglycerol (2-AG) are produced endogenously and function as a partial agonist and a full agonist, respectively, at both receptors (Zou and Kumar, 2018). Δ⁹-THC, the principal psychoactive constituent of cannabis, is a partial agonist (Pertwee, 2008). Synthetic cannabinoid receptor agonists (SCRAs) such as those found in illicit “K2/Spice” products, are full or higher-efficacy agonists (Atwood et al., 2010). Among antagonists, inverse agonists and neutral antagonists have been studied for different therapeutic applications, with neutral antagonists generally preferred for neuropsychiatric indications owing to the adverse psychiatric profile associated with inverse agonism (Gueye et al., 2016).

Although the pharmacopeia of CB1 and CB2 ligands has been built from collective information across numerous pharmacological studies, those prior characterizations have typically examined structurally narrow subsets of compounds. Furthermore, those individual studies often used differing cell systems and assay conditions, making efficacy difficult to compare across ligands (Ibsen et al., 2019). Evaluation of ligands with this efficacy and structural breadth has not been reported within a single well-controlled assay system at both CB1 and CB2. Furthermore, many prior readouts, cAMP inhibition chief among them, are indirect and heavily amplified, so measured efficacies and potencies may be conflated with receptor reserve and assay amplification (Wang et al., 2004). A complete account requires resolving how ligands engage the two major transducer pathways, Gαi1 and β-arrestin 2, because the balance between them can drive distinct physiological, therapeutic, and abuse-relevant outcomes (Ibsen et al., 2017).

Here we address these limitations using bioluminescence resonance energy transfer (BRET) assays. By measuring the direct coupling between receptor and Gα, or between receptor and β-arrestin, in real time within intact cells, BRET reports engagement at the level of the receptor transducer complex rather than an amplified downstream second messenger (Namkung et al., 2016; Olsen et al., 2020). Using the same cultured cells, assay format, and the reference ligand across the entire panel and for both CB1 and CB2, we determine dose-response relationships for Gαi1 engagement and β-arrestin 2 recruitment across compounds spanning the full efficacy spectrum: endocannabinoid analogues, Δ⁹-THC, SCRAs, and antagonists, at both CB1 and CB2 under uniform conditions. The result is a systematic efficacy and potency map across a structurally and functionally diverse panel at both receptors on two transducers, enabling the direct cross-compound comparison that fragmented literature cannot support. In total, the panel comprised 22 cannabinoid ligands across four structural classes, each profiled at CB1 and CB2 on both the Gαi1 engagement and β-arrestin 2 recruitment assays.

## Methods

### Cell Culture and Transfection

Human embryonic kidney 293T (HEK-293T) cells were grown in Dulbecco’s modified Eagle’s medium (DMEM, Corning) containing 10% fetal bovine serum, 2 mM l-glutamine (Corning), and 1% penicillin–streptomycin (Corning). Cultures were kept on 10 cm plates at 37 °C in 5% CO2. For each experiment, 3 × 10⁶ cells were plated per 10 cm dish and allowed to become confluent over 24 h before transfection. After overnight transfection (16 h), medium was exchanged for fresh DMEM, and assays were carried out 48 h post-transfection.

HEK-293T cells were transiently transfected with cDNA (10–15 μg per assay) and linear polyethylenimine hydrochloride (PEI Max, Polysciences) at a 1:2 ratio (30 μg). Plasmid mass and transfection ratio were optimized and kept consistent across BRET assays. The reagent–plasmid mixture was applied dropwise to the medium and left to incubate overnight.

### Bioluminescence Resonance Energy Transfer (BRET) Assays

Two BRET assays were performed in this study, measuring Gαi1 engagement and β-arrestin 2 recruitment. For the Gαi1 engagement assay, the resonance energy transfer (RET) pair comprised a Renilla luciferase 8 (RLuc)–fused receptor and Venus-fused Gαi1. To avoid depleting endogenous Gβγ, Gβ1 and Gγ2 constructs were also cotransfected. For β-arrestin 2 recruitment, the RET pair consisted of an RLuc-fused receptor and Venus-fused β-arrestin 2. GRK2 constructs were cotransfected to increase receptor phosphorylation and thereby facilitate β-arrestin 2 recruitment.

At 48 h post-transfection, cells were rinsed with PBS, detached with 5 mM EDTA (Fisher Scientific), harvested, and resuspended in PBS supplemented with 0.1% glucose, 0.1% BSA, and 200 μM sodium bisulfite. The suspension was dispensed into white U-bottom 96-well plates, and ligands were applied as either agonists or antagonists. Agonists were added in triplicate as 10-fold serial dilutions. For antagonist conditions, cells were preincubated with 1 μM antagonist for 10 min before the agonist CP55940 was added. For antagonist measurements, antagonists were added alone as 10-fold serial dilutions in the absence of agonist. For competitive antagonism, antagonists were titrated as 10-fold serial dilutions in the presence of a fixed concentration of CP55,940. Cells were incubated with the luciferase substrate coelenterazine H (5 μM) for 2.5 min before agonist addition. G protein or β-arrestin 2 engagement and dissociation were quantified from BRET ratios (530/480 nm) measured between the RLuc- and Venus-tagged proteins. All conditions were run in three to five biological replicates.

## Compounds

CP55,940 and all other cannabinoid ligands were given by the Makriyannis Laboratory, obtained from Cayman Chemical (Ann Arbor, MI) or the NIDA Drug Supply Program (Rockville, MD) and dissolved in DMSO at 10 mM.

### Statistical Analysis

BRET ratios (530/480 nm) were used to construct dose response curves and compared by one-way ANOVA in GraphPad Prism v10. All data were normalized to the maximal and minimal BRET signal ratios of the reference agonist CP55940. Specifically, Emax values for every ligand were scaled to the maximal CP55940 response within each assay (e.g., Gαi1 engagement or β-arrestin 2 recruitment). This approach permits direct comparison of relative efficacy across ligands, both within and between assays. Emax and pEC50 values were obtained by nonlinear regression of the pooled data.

Differences in Emax and pEC50 relative to CP55,940 were assessed by one-way ANOVA with Dunnett’s multiple-comparisons test, and antagonist parameters were compared to AM251 by the same method. Differences between CB1 and CB2 (for a given transducer) and between Gαi1 engagement and β-arrestin 2 recruitment (for a given receptor) were assessed within each structural group by two-way ANOVA with Šidak’s multiple-comparisons test. Signaling bias was evaluated from extrapolated Emax and EC50 values: for each transducer the log(Emax/EC50) of each ligand was referenced to that of CP55,940 to obtain Δlog(Emax/EC50), and the ΔΔlog(Emax/EC50) for Gαi1 engagement versus β-arrestin 2 recruitment was calculated, with a value greater than 1 or less than −1 taken to indicate at least a 10-fold transducer preference. For dose response curves that did not reach a plateau, Emax and pEC50 were recorded as not determined (N.D.). Curve colors in all dose–response figures were selected from colorblind-safe ColorBrewer palettes (colorbrewer2.org) to maximize visual contrast among the ligands shown within each panel.

## Results

To characterize cannabinoid receptor ligands across all efficacies, we used a structurally diverse panel of compounds and profiled them at both CB1 and CB2 receptors. The panel was organized structurally into four classes: synthetic cannabinoid receptor agonists (SCRAs), Δ⁹-tetrahydrocannabinol (Δ⁹-THC) analogues, endocannabinoid analogues, and antagonists/inverse agonists. These ligands span the range from high-efficacy agonists to inverse agonists within a uniform BRET assay, pairing well-established reference ligands within each class with AM compounds and

SCRAs. Agonist responses were normalized to the reference full agonist CP55,940 (100% Emax), which was selected for its stability and high potency.

The SCRA ligand group consisted of the “third-generation” indazole- and indole-3-carboxamide agonists MDMB-4en-PINACA, MDMB-FUB, and 5F-MDMB-PICA, the naphthoylindole AM2201, and the aminoalkylindole WIN55,212-2. The three carboxamide SCRAs are prominent drugs of abuse that exceed the efficacy of classical reference agonists in G-protein and β-arrestin readouts and can be classified as high-efficacy agonists (Ibsen et al., 2017; Krotulski et al., 2021). AM2201 and WIN55,212-2 were additionally included as established reference agonists. AM2201 is a widely used standard, particularly in electrophysiology, and WIN55,212-2 is a nonselective full agonist routinely used to normalize cannabinoid binding and functional activity, and is also the closest structural relative to the carboxamide SCRAs (Banister et al., 2015; Felder et al., 1995; Ibsen et al., 2017).

The Δ⁹-THC analogue group consisted of Δ⁹-THC, a low-efficacy partial agonist at both CB1 and CB2 receptors, as well as the high-affinity, CB1-selective agonist AM2389, and AM2302, AM8936, and AM11101 (Ibsen et al., 2017; Järbe et al., 2012; Nikas et al., 2010). Of these, AM8936 has been described as a potent, efficacious CB1 agonist and AM11101 as a CB1 partial agonist (Malamas et al., 2018; Papanastasiou et al., 2022).

The endocannabinoid analogue group comprised the endogenous ligands anandamide (AEA) and 2-arachidonoylglycerol (2-AG) alongside the metabolically stabilized analogues AM356 (methanandamide), AM8125, AM12814, and AMG315. AEA is a partial agonist at CB1 with low intrinsic activity at CB2, whereas 2-AG behaves as a full agonist at both receptors (Abadji et al., 1994; Gonsiorek et al., 2000). AM356 is a widely used, metabolically stable AEA reference analogue, while AM8125, AM12814, and AMG315 are ligands described in single originating reports.

The antagonist/inverse-agonist group included rimonabant (SR141716A) and AM251, the two most widely used reference CB1 inverse agonists, together with AM6538, a competitive antagonist used to determine the first CB1 crystal structure, and the neutral antagonists AM4113, AM6545, and AM6527. Among the neutral antagonists, AM4113 is the classic CB1 neutral antagonist and AM6545 is a widely used peripherally restricted neutral antagonist, whereas AM6527 is an orally active neutral antagonist (Cluny et al., 2010; Sink et al., 2009, 2008).

Because these prior efficacy classifications derive largely from cAMP-inhibition and [³⁵S]GTPγS G-protein assays and, for several SCRAs, from β-arrestin-recruitment assays, and were often normalized to reference agonists other than CP55,940, they are not directly comparable across studies. We therefore evaluated the efficacy and potency of all ligands at CB1 and CB2 using bioluminescence resonance energy transfer (BRET)-based biosensors to monitor (1) receptor interaction with the Gαi1 subunit (Gαi1 engagement) and (2) receptor recruitment of β-arrestin 2 (β-arrestin 2 recruitment). Profiling both readouts at each receptor produced four receptor transducer measurements: CB1 Gαi1 engagement, CB1 β-arrestin 2 recruitment, CB2 Gαi1 engagement, and CB2 β-arrestin 2 recruitment. These biosensors report proximal receptor transducer coupling directly and are insensitive to off-target receptor activation, enabling a direct comparison of Gαi1 and β-arrestin 2 engagement across the efficacy spectrum.

### At the CB1 Gαi1 engagement assay, SCRAs and several AM compounds display high efficacy

At CB1, SCRAs were the most efficacious agonists in the Gαi1 engagement assay, each exceeding the CP55,940 maximum (p<0.0001). MDMB-4en-PINACA produced the largest response (Emax 167.1 ± 3.7%), followed by 5F-MDMB-PICA (154.1 ± 1.8%) and MDMB-FUB (140.6 ± 2.5%). Of these, MDMB-FUB was the most potent (pEC50 8.42 ± 0.06), comparable to that of CP55,940 (**Figure 1A**, **Table 1**). The two aminoalkylindole reference agonists were both full agonists, but only AM2201 matched the CP55,940 Emax (110.2 ± 1.5%, not significantly different). WIN55,212-2 modestly exceeded it (116.3 ± 4.3%, p<0.0001), while remaining below the higher-efficacy SCRAs. They diverged markedly in potency: AM2201 was highly potent (pEC50 8.51 ± 0.05), whereas WIN55,212-2 was right-shifted by approximately two log units (pEC50 6.25 ± 0.07).

**Figure 1.**
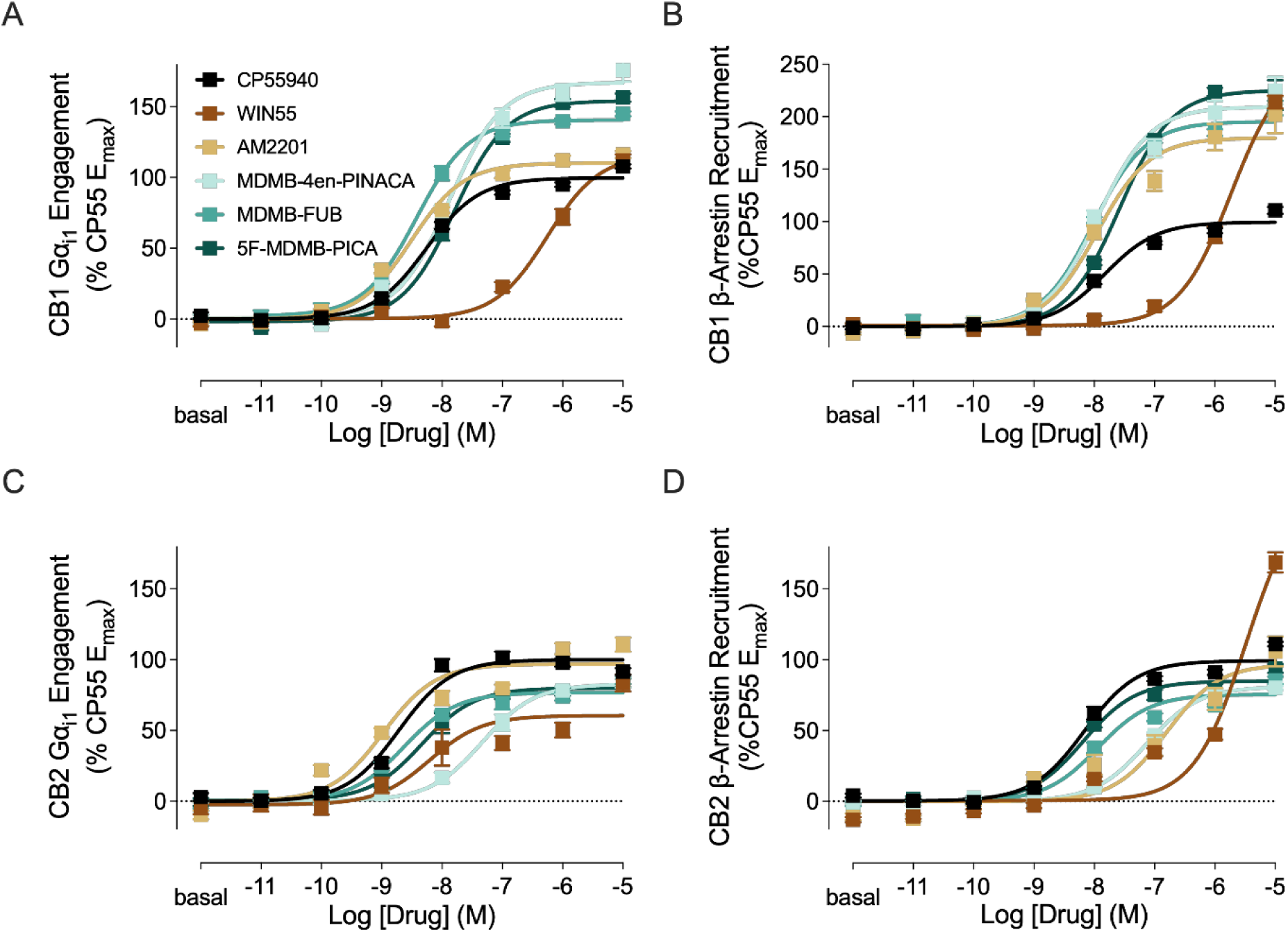
Dose response of high-efficacy synthetic cannabinoid receptor agonists at CB1 and CB2 receptors. BRET concentration–response curves for CP55940, WIN55,212-2, AM2201, MDMB-4en-PINACA, MDMB-FUB, and 5F-MDMB-PICA measuring (**A**) CB1 Gαi1 engagement, (**B**) CB1 β-arrestin 2 recruitment, (**C**) CB2 Gαi1 engagement, and (**D**) CB2 β-arrestin 2 recruitment. Data are presented as mean ± SEM (n = 3-5).

**Table 1.**
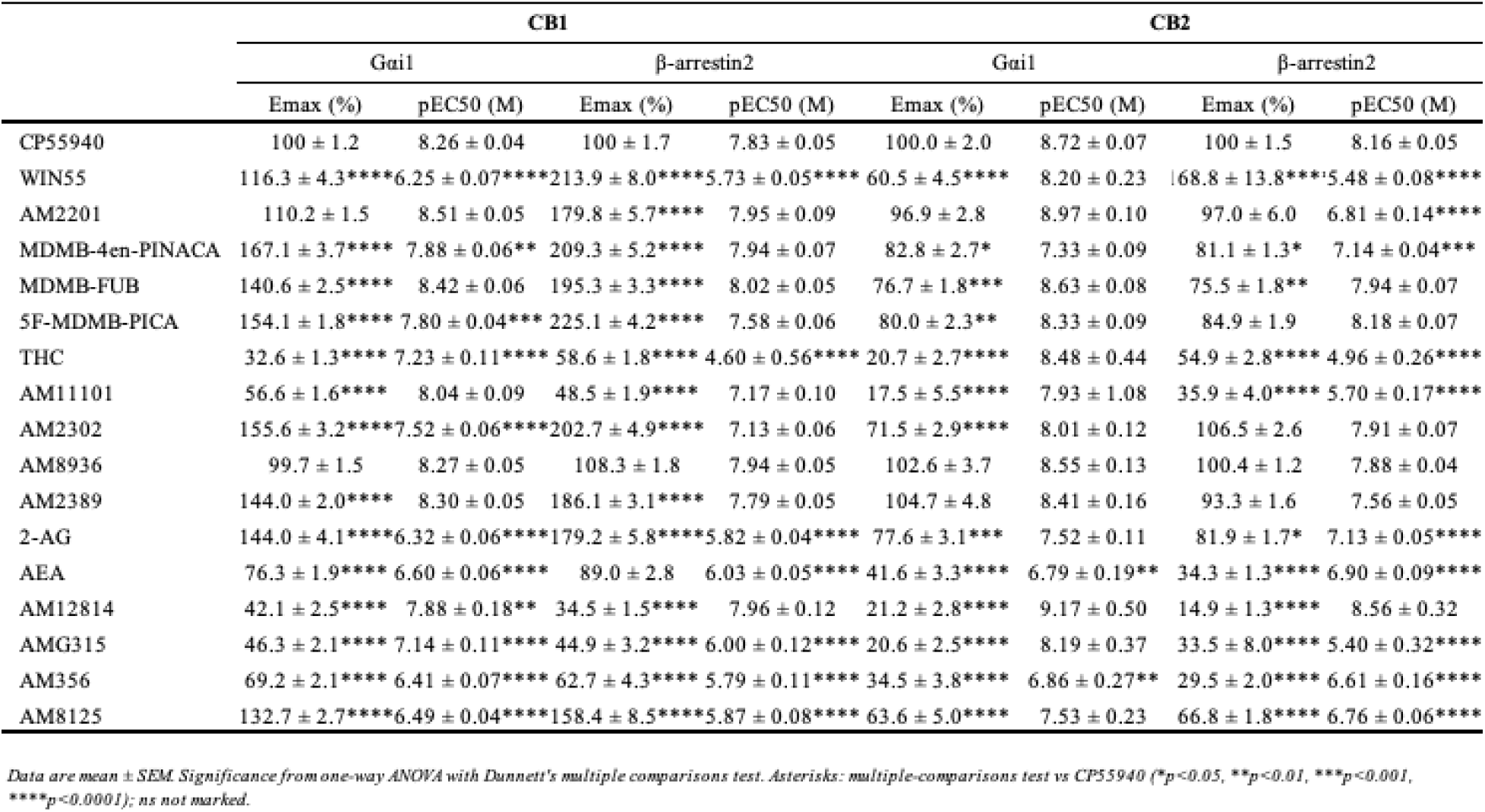
Efficacy and potency of all compounds.

**Table 2.** Bias factors against CP55940 as reference.

| | $\Delta\log(E_{max}/EC_{50})$ | |
| --- | --- | --- |
|  | CB1R | CB2R |
| WIN55 | N.D. | N.D. |
| AM2201 | -0.08 | 1.61 |
| MDMB-4en-PINACA | -0.59 | -0.35 |
| MDMB-FUB | -0.17 | 0.14 |
| 5F-MDMB-PICA | -0.38 | -0.43 |
| AM2302 | -0.16 | -0.63 |
| AM8936 | -0.13 | 0.12 |
| AM2389 | -0.04 | 0.34 |
| AM11101 | 0.50 | N.D. |
| THC | N.D. | N.D. |
| 2-AG (AM Comp) | 0.15 | -0.18 |
| AEA (AM Comp) | N.D. | -0.59 |
| AM12814 | -0.42 | 0.21 |
| AMG315 | 0.72 | N.D. |
| AM356 | N.D. | -0.24 |
| AM8125 | N.D. | 0.19 |

The Δ⁹-THC analogues had the widest efficacy range in the Gαi1 engagement assay. Δ⁹-THC was a low-efficacy partial agonist (Emax 32.6 ± 1.3%) and AM11101 was likewise partial (56.6 ± 1.6%), whereas AM2302 and AM2389 were highly efficacious (155.6 ± 3.2% and 144.0 ± 2.0%, respectively) and AM8936 tracked CP55,940 closely in both efficacy and potency (99.7 ± 1.5%; pEC50 8.27 ± 0.05; not significantly different from CP55,940) (**Figure 2A**, **Table 1**). The low efficacy of Δ⁹-THC in this assay as well as its low CB2 Gαi1 response (20.7 ± 2.7%) is consistent with prior [³⁵S]GTPγS data that showed Δ⁹-THC as a partial agonist that reaches only ∼20–50% of the CP55,940 response (Pertwee, 2008).

**Figure 2.**
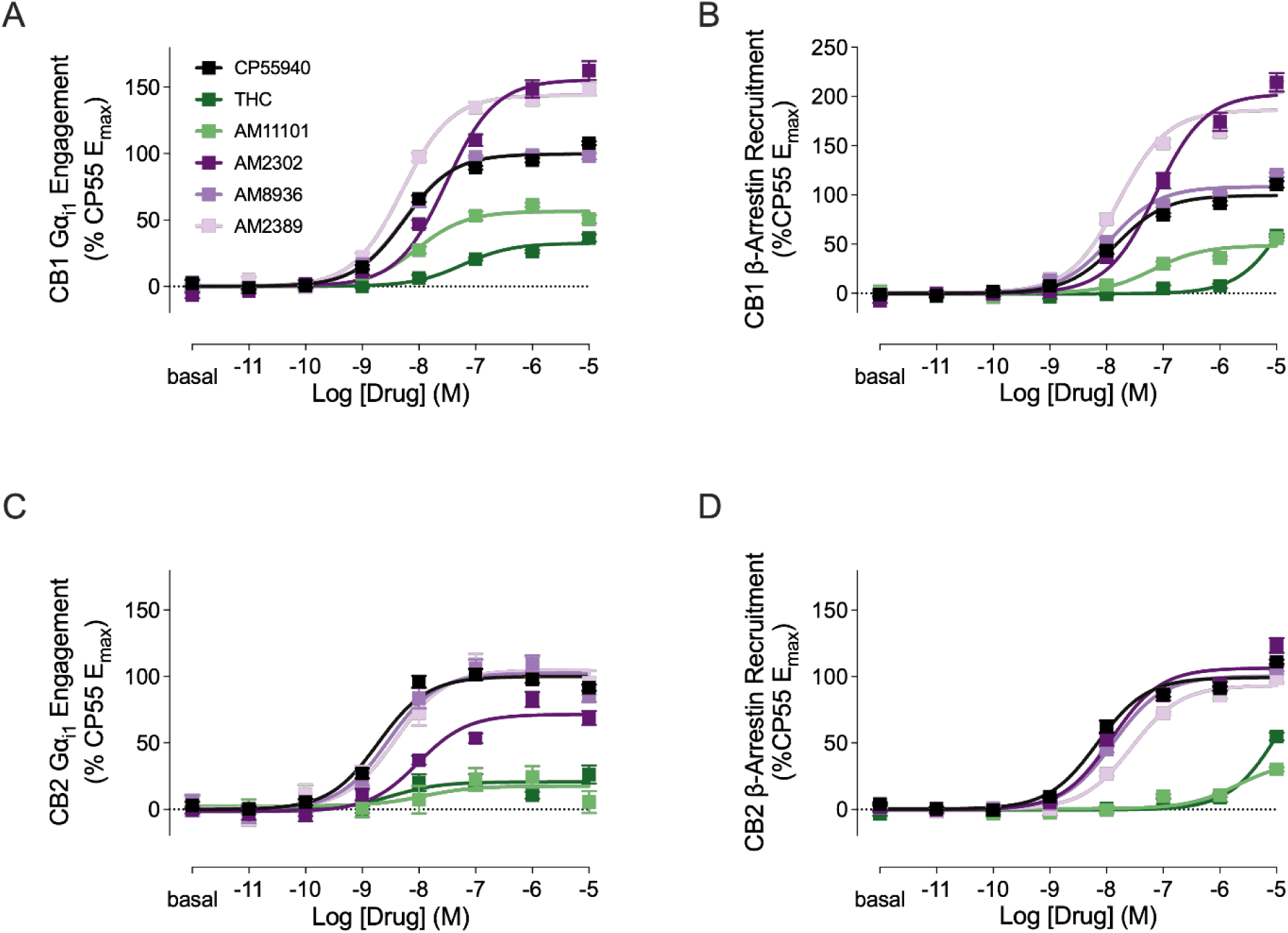
Dose response of Δ⁹-THC and its analogues at CB1 and CB2 receptors. BRET concentration–response curves for CP55940, Δ⁹-THC, AM11101, AM2302, AM8936, and AM2389 measuring (**A**) CB1 Gαi1 engagement, (**B**) CB1 β-arrestin 2 recruitment, (**C**) CB2 Gαi1 engagement, and (**D**) CB2 β-arrestin 2 recruitment. Data are presented as mean ± SEM (n = 3–5).

Among the endocannabinoid analogues, 2-AG, AEA, AM8125, and AM356 were relatively low in potency (pEC50 6.32–6.60), with 2-AG (Emax 144.0 ± 4.1%) and AM8125 (132.7 ± 2.7%) showing high efficacy while AEA (76.3 ± 1.9%) and AM356 (69.2 ± 2.1%) remained partial. AM12814 and AMG315 were more potent (pEC50 7.88 ± 0.18 and 7.14 ± 0.11) but low in efficacy (42.1 ± 2.5% and 46.3 ± 2.1%) (**Figure 3A**, **Table 1**). Overall, at CB1 Gαi1 engagement, high efficacy was common among the SCRAs and several Δ⁹-THC analogues.

**Figure 3.**
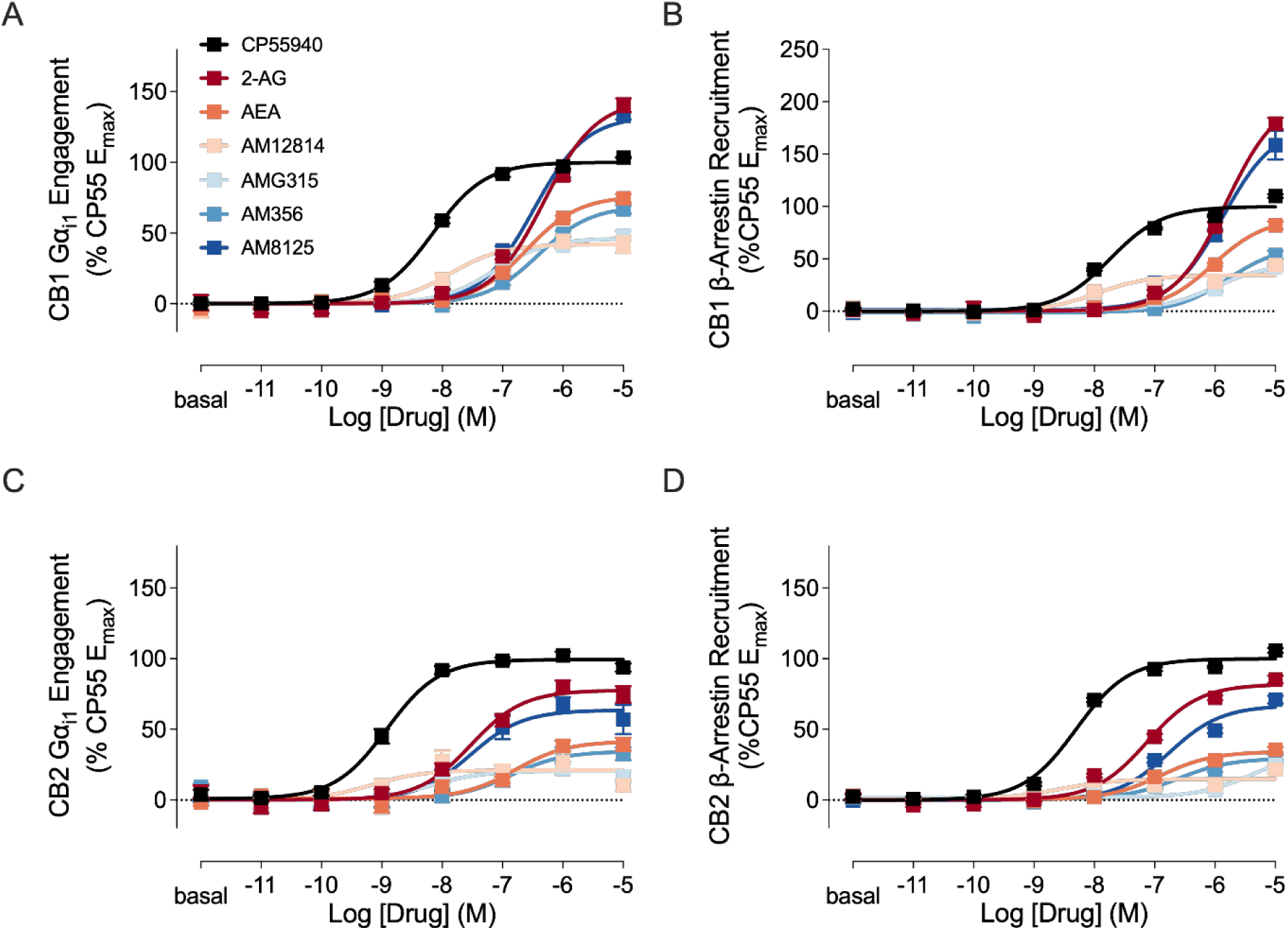
Dose response of endocannabinoids and their analogues at CB1 and CB2 receptors. BRET concentration–response curves for CP55940, 2-AG, AEA, AM12814, AMG315, AM356, and AM8125 measuring (**A**) CB1 Gαi1 engagement, (**B**) CB1 β-arrestin 2 recruitment, (**C**) CB2 Gαi1 engagement, and (**D**) CB2 β-arrestin 2 recruitment. Data are presented as mean ± SEM (n = 3–5).

### High-efficacy agonists recruit β-arrestin 2 above CP55,940 at CB1

In the CB1 β-arrestin-2 recruitment assay, efficacy was substantially higher than in the Gαi1 engagement assay, and the Emax differences from CP55,940 were significant for every compound except AM8936 and AEA (**Table 1**). The SCRAs highly recruited β-arrestin 2 to 195–225%, with 5F-MDMB-PICA giving the highest efficacy in the entire assay (225.1 ± 4.2%) while remaining potent (pEC50 7.58–8.02). AM2201 similarly recruited (179.8 ± 5.7%, p<0.0001) with high potency (pEC50 7.95 ± 0.09), whereas WIN55,212-2 also recruited efficaciously (213.9 ± 8.0%, p<0.0001) but at low potency (pEC50 5.73 ± 0.05) (**Figure 1B**, **Table 1**).

Δ⁹-THC recruited weakly and with the lowest potency in the panel (Emax 58.6 ± 1.8%; pEC50 4.60 ± 0.56), and AM11101 was a partial recruiter (48.5 ± 1.9%), both significantly below CP55,940 (p<0.0001). In contrast, AM2302 (202.7 ± 4.9%) and AM2389 (186.1 ± 3.1%) recruited strongly (both p<0.0001). AM8936 alone recruited at the CP55,940 level (108.3 ± 1.8%, not significantly different) (**Figure 2B**, **Table 1**).

Within the endocannabinoid analogues, 2-AG (Emax 179.2 ± 5.8%) and AM8125 (158.4 ± 8.5%) recruited significantly (both p<0.0001) despite low potency (pEC50 5.82 ± 0.04 and 5.87 ± 0.08), whereas AM12814 (34.5 ± 1.5%), AM356 (62.7 ± 4.3%), and AMG315 (44.9 ± 3.2%) recruited significantly below CP55,940 (p<0.0001). AM12814, though the least efficacious, was the most potent of the group (pEC50 7.96 ± 0.12), and AEA alone was indistinguishable from CP55,940 (89.0 ± 2.8%) (**Figure 3B**, **Table 1**). β-arrestin-2 recruitment at CB1 was therefore pronounced and significant. The SCRAs, AM2201, 2-AG, and AM8125 all exceeded CP55,940 at p<0.0001, whereas the endocannabinoids, WIN55,212-2, and Δ⁹-THC were sharply right-shifted in potency (pEC50 4.6–6.0).

### CB2 Gαi1 efficacies are low relative to CB1, largely without high efficacy

At the CB2 Gαi1 engagement assay, SCRAs that had been highly efficacious at CB1 were reduced to partial agonists (Emax 76.7–82.8%): MDMB-4en-PINACA (82.8 ± 2.7%), 5F-MDMB-PICA (80.0 ± 2.3%), and MDMB-FUB (76.7 ± 1.8%), but still the most potent (pEC50 8.63 ± 0.08) (**Figure 1C**, **Table 1**). This CB1 to CB2 efficacy reduction is notable compared to prior work: Banister et al. reported MDMB-FUB as a full agonist at both receptors (Emax 108% at CB1 and 101% at CB2, normalized to CP55,940) with a modest CB1 preference (Banister et al., 2016). Here the CB1 preference was reproduced but was considerably more pronounced, with CB1 Gαi1 high efficacy (140.6 ± 2.5%) giving way to only partial CB2 Gαi1 efficacy (76.7 ± 1.8%). AM2201 was the exception among the SCRAs, with the highest efficacy in at CB2 Gαi1 (96.9 ± 2.8%) and a low fitted EC50 (pEC50 8.97 ± 0.10). WIN55,212-2 showed the lowest SCRA efficacy at CB2 Gαi1 (60.5 ± 4.5%), yet in contrast to its low CB1 potency, was potent here (pEC50 8.20 ± 0.23). Prior work classifies WIN55,212-2 as an essentially non-selective full agonist (CB2 Ki 3.3 nM; CB1 Ki up to 62.3 nM at human recombinant CB1), and the greater CB2 than CB1 Gαi1 potency observed here is consistent with that more potent CB2 affinity, although the reduction to partial efficacy at CB2 is different from the earlier full-agonist classification (Felder et al., 1995).

The Δ⁹-THC analogues behaved differently at CB2 Gαi1: AM2389 (Emax 104.7 ± 4.8%) and AM8936 (102.6 ± 3.7%) were full agonists matched to CP55,940, whereas Δ⁹-THC (20.7 ± 2.7%) and AM11101 (17.5 ± 5.5%) were minimally efficacious, and AM2302 was intermediate (71.5 ± 2.9%) (**Figure 2C**, **Table 1**).

The endocannabinoid analogues had uniformly low efficacies at CB2 Gαi1 (Emax 20.6– 77.6%). 2-AG was the most efficacious of the group (77.6 ± 3.1%). Notably, although 2-AG has been reported as an endogenous full agonist at both receptors ([³⁵S]GTPγS EC50 38.9 nM at human CB2), it behaved as only a partial agonist at CB2 Gαi1 here, consistent with its high efficacy at CB1 but below the prior full-agonist classification at CB2. AM12814 was the most potent compound in the dataset overall (pEC50 9.17 ± 0.50) despite minimal efficacy (21.2 ± 2.8%) (**Figure 3C**, **Table 1**). In sum, CB2 Gαi1 efficacies fell sharply relative to CB1: high efficacies largely disappeared, most agonists were partial, and potencies were generally high (several pEC50 > 8).

### At CB2 β-Arrestin-2, most ligands recruit at or below CP55,940

In the CB2 β-arrestin-2 recruitment assay, the SCRAs were partial recruiters (Emax ∼75–85%): MDMB-4en-PINACA (81.1 ± 1.3%) and MDMB-FUB (75.5 ± 1.8%) both recruited significantly below CP55,940 (p<0.05 and p<0.01, respectively), whereas 5F-MDMB-PICA (84.9 ± 1.9%, the most potent at pEC50 8.18 ± 0.07) did not differ significantly. AM2201 recruited to the CP55,940 level (97.0 ± 6.0%, not significantly different) but with right-shifted potency (pEC50 6.81 ± 0.14), while WIN55,212-2 was again the outlier, recruiting efficaciously (168.8 ± 13.8%, p<0.0001) at the lowest potency in the panel (pEC50 5.48 ± 0.08) (**Figure 1D**, **Table 1**).

Among the Δ⁹-THC analogues, AM2302 (Emax 106.5 ± 2.6%), AM8936 (100.4 ± 1.2%), and AM2389 (93.3 ± 1.6%) recruited at or near the CP55,940 level (all not significantly different), whereas Δ⁹-THC (54.9 ± 2.8%) and AM11101 (35.9 ± 4.0%) were significantly weaker (both p<0.0001) (**Figure 2D**, **Table 1**). The endocannabinoid analogues were predominantly low-efficacy partial recruiters that fell significantly below CP55,940. 2-AG was the least reduced, at 81.9 ± 1.7% (p<0.05), and AEA, AM8125, AM356, AMG315, and AM12814 were all significant at p<0.0001 with AM12814 the lowest in the entire panel (14.9 ± 1.3%) yet still potent (pEC50 8.56 ± 0.32) (**Figure 3D**, **Table 1**). Thus, at CB2 β-arrestin-2, most compounds recruited significantly below the CP55,940 maximum, with WIN55,212-2 the sole marked enhanced recruiter and the AM Δ⁹-THC analogues AM2302, AM2389, and AM8936 recruiting fully.

### Inverse Agonists Suppress Constitutive CB1 Activity More Than Neutral Antagonists

The six ligands in the CB1 antagonist group comprise the two established inverse agonists AM251 and rimonabant, together with AM6538, a structural analogue of rimonabant and competitive antagonist that behaves as an inverse agonist, as well as three neutral antagonists, AM4113, AM6545, and AM6527 (Cluny et al., 2010; Hua et al., 2016; Lan et al., 1999; Landsman et al., 1997; Rinaldi-Carmona et al., 1994; Sink et al., 2009, 2008). We first assayed them alone against constitutive CB1 Gαi1 activity to measure its intrinsic inhibition efficacy (**Figure 4A**, **Table 3**). The inverse agonists reduced basal signaling most (−19 to −22% of the CP55,940 window) and at high potency (pEC50 7.8–8.6). The neutral antagonists AM4113 (−12.8%) and AM6545 (−12.3%), and AM6527 (−127%) reduced constitutive activity significantly less than AM251 (all p<0.01), consistent with a weaker or absent inverse agonist signature, and were also significantly less potent than AM251 (pEC50 6.9-7.1, p<0.001).

**Figure 4.**
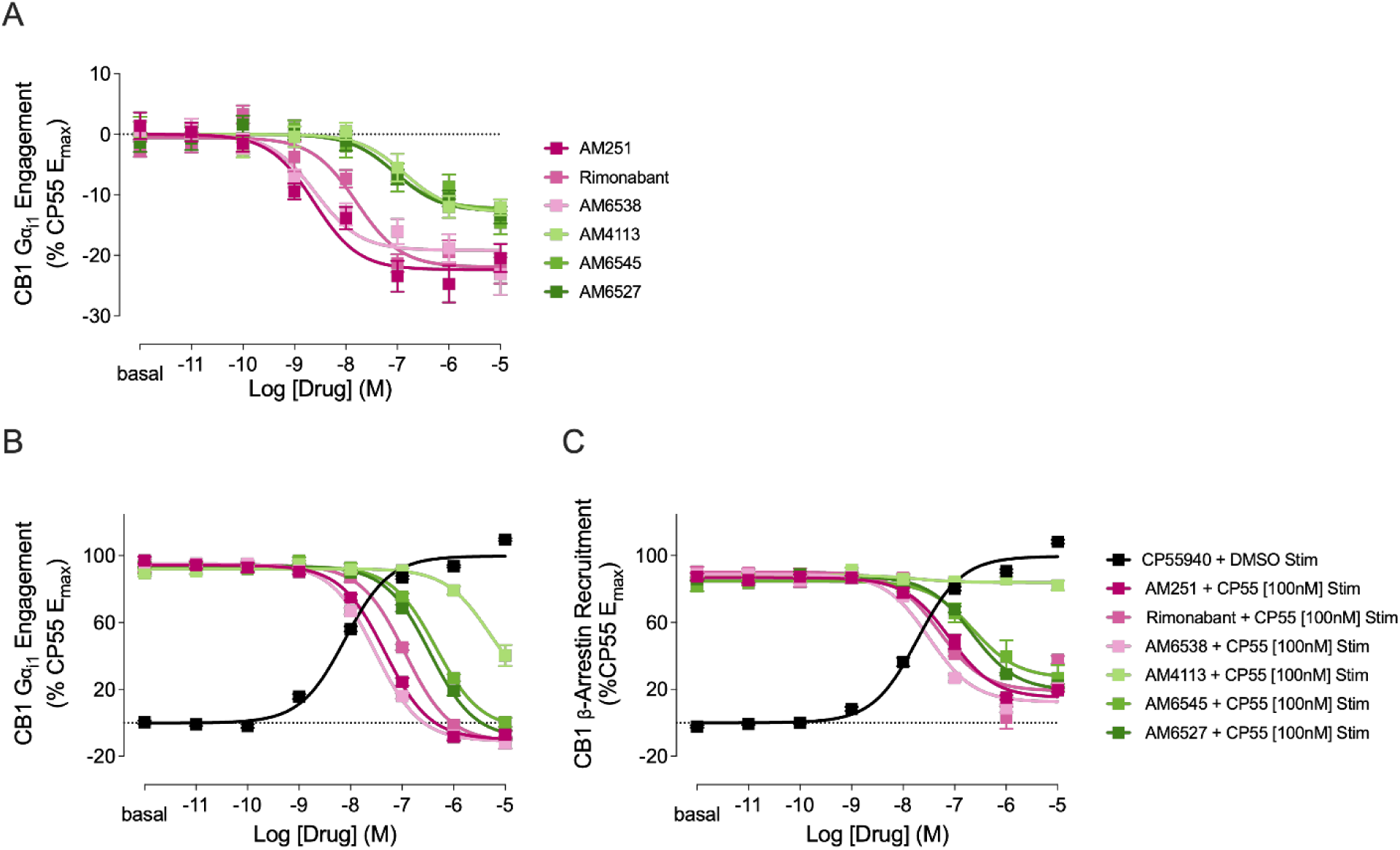
Dose response of cannabinoid antagonists at CB1. BRET concentration–response curves at CB1 for AM251, Rimonabant, AM6538, AM4113, AM6545, and AM6527 measuring (**A**) Gαi1 engagement by each compound alone. (**B**) Gαi1 engagement and (**C**) β-arrestin 2 recruitment for a CP55940 dose response in the presence of vehicle (DMSO), overlaid with each antagonist titrated in the presence of a fixed 100 nM CP55940. Data are presented as mean ± SEM (n = 3–5).

**Table 3.** Antagonist activity on basal condition.

|  | **CBI** |  |
| --- | --- | --- |
|  | Gai1 |  |
|  | Emax (%) | pEC50 (M) |
| AM251 | -22.4 ± 1.27 | 8.62 ± 0.19 |
| Rimonabant | -21.4 ± 1.25 | 7.79 ± 0.17 |
| AM6538 | -19.2 ± 1.22 | 8.60 ± 0.22 |
| AM4113 | -12.8 ± 1.33** | 6.87 ± 0.24**** |
| AM6545 | -12.3 ± 1.49** | 7.07 ± 0.29**** |
| AM6527 | -12.7 ± 1.21** | 7.04 ± 0.23*** |

Data are mean ± SEM; Emax normalized to CP55,940 = 100%. Ligands assayed alone. Significance from one-way ANOVA with Dunnett's multiple comparisons test vs AM251 (\*p ≤ 0.05, \*\*p ≤ 0.01, \*\*\*p ≤ 0.001, \*\*\*\*p ≤ 0.0001); ns not marked

Against a saturating stimulation with 100 nM CP55,940 (**Figure 4B,C**; **Table 4**), the inverse agonists drove the Gαi1 response below the CP55,940-defined baseline (Imax −10 to −11%), reflecting inverse agonism on top of full blockade, and were the most potent blockers (pIC50 6.9–7.5). AM4113 was the exception: its Gαi1 blockade was incomplete (Imax 14.1%, significantly different from AM251, p<0.05) and the least potent (pIC50 5.30), and in β-arrestin-2 it barely blocked at all (Imax 83.9%, the curve remaining near the CP55,940 level), whereas the other five reduced β-arrestin-2 recruitment to 12–27%. The two other neutral antagonists, AM6545 and AM6527, blocked both pathways effectively.

**Table 4.** Antagonist activity with pre-stimulation of 100 nM CP55940.

|  | **CBI** |  |  |  |
| --- | --- | --- | --- | --- |
|  | Gai1 |  | β-arrestin2 |  |
|  | Imax (%) | pIC50 (M) | Imax (%) | pIC50 (M) |
| AM251 | -10.0 ± 1.67 | 7.33 ± 0.04 | 14.9 ± 1.85 | 7.10 ± 0.06 |
| Rimonabant | -11.3 ± 1.64 | 6.94 ± 0.04 | 19.1 ± 5.39 | 7.30 ± 0.20 |
| AM6538 | -10.9 ± 1.46 | 7.52 ± 0.04 | 12.4 ± 1.63 | 7.49 ± 0.06 |
| AM4113 | 14.1 ± 13.72* | 5.30 ± 0.20 | 83.9 ± 1.20 | 7.77 ± 0.77 |
| AM6545 | -3.8 ± 2.44 | 6.34 ± 0.05 | 26.8 ± 4.97 | 6.64 ± 0.20 |
| AM6527 | -9.3 ± 2.35 | 6.47 ± 0.05 | 19.3 ± 1.35 | 6.66 ± 0.05 |

Data are mean ± SEM; Imax normalized to CP55,940 = 100%. Antagonists assayed against a fixed concentration of CP55,940. Significance from one-way ANOVA with Dunnett's multiple comparisons test vs AM251 (\*p ≤ 0.05, \*\*p ≤ 0.01, \*\*\*p ≤ 0.001, \*\*\*\*p ≤ 0.0001); ns not marked. N.D., not determined (low-potency curve); Imax shown is the effect at the maximum concentration tested (10 μM).

## Discussion

The most consistent feature of the panel was a loss of agonist efficacy from CB1 to CB2 (**Supplementary Tables S1–S3**). Carboxamide SCRAs that had high efficacies at CB1 (140–167% of CP55,940) fell to partial agonism at CB2 (77–83%); the Δ⁹-THC analogues with high efficacy AM2302 and AM2389 dropped in parallel; and every endocannabinoid was less efficacious at CB2, with the anandamide analogues AM12814 and AMG315 declining to near-inactivity. Only two agonists remained the same in the Gαi1 engagement assay. AM8936, a full agonist at both receptors, and AM2201, which appeared to remain close to a full agonist at CB2. Potency remained unchanged or even increased at CB2 (uniformly for the endocannabinoid analogues, variably for the SCRAs).

A second receptor dependent pattern occurred between the two transducers β-arrestin 2 and Gαi1. At CB1, the high-efficacy agonists of every class recruited β-arrestin 2 to efficacies well above both their own Gαi1 responses and CP55,940, which included the carboxamide SCRAs, AM2201, and WIN55,212-2, together with Δ⁹-THC, AM2302, and AM2389 and the endocannabinoids 2-AG and AM8125. This extensive recruitment was specific to CB1 **(Supplementary Tables S1–S3)**. It should be noted that the high-efficacy behavior observed for many ligands in the CB1-β-arrestin 2 assay may reflect the lower relative efficacy of CP55,940 in this assay rather than unusually high ligand efficacy. Regardless, the relative differences among ligands remain unaffected by the choice of reference. At CB2, most ligands showed comparable efficacies at β-arrestin-2 and Gαi1, with only WIN55,212-2, Δ⁹-THC, AM2302, AM11101, and AMG315 exhibiting greater efficacy at β-arrestin-2 than at Gαi1.

Three ligands stood out against the trends mentioned. AM8936 was a balanced agonist throughout the panel, a full agonist and CP55,940-like at both receptors and both transducers. WIN55,212-2 sat at the opposite extreme: it retained high efficacy at β-arrestin-2 at both receptors and maintained an unusually low Gαi1 potency. AM2201 showed high efficacies at CB2 and was the only compound in the panel to show a signaling bias, while the SCRA carboxamides showed partial agonism at CB2.

Bias analysis reinforced this picture. Bias factors were calculated from the extrapolated Emax and EC50 values (**Table 2**). For each transducer we determined the log(Emax/EC50) of every ligand, subtracted the log(Emax/EC50) of CP55,940 to obtain the Δlog(Emax/EC50), and then subtracted the Δlog(Emax/EC50) values of the two transducers to obtain the ΔΔlog(Emax/EC50) for Gαi1 versus β-arrestin-2, with positive values indicating a Gαi1 preference and negative values a β-arrestin-2 preference.

Because these are logarithmic values, a bias factor greater than 1 or less than −1 reflects a ≥10-fold difference in transducer selectivity, thus, almost all ligands with a well-defined dose response curve fell within the category of balanced agonist. At CB1, no determinable ΔΔlog fell outside this threshold; at CB2, the sole biased ligand was AM2201, with a modest Gαi1 preference (ΔΔlog(Emax/EC50) = +1.61). Several compounds could not be assigned a bias factor because their β-arrestin-2 curves did not plateau: WIN55,212-2, Δ⁹-THC, AEA, AM356, and AM8125 at CB1, and WIN55,212-2, AM11101, Δ⁹-THC, and AMG315 at CB2, so those assignments remain inconclusive. The broader implication is that the marked efficacy and receptor-subtype differences across the panel do not reflect pathway-selective signaling with the single exception of AM2201 at CB2, the ligands differ in how strongly they activate each receptor, not in which transducer they prefer.

The antagonist panel behaved consistently with previously established categorization of neutral agonists and inverse agonists. Against constitutive CB1 Gαi1 activity, the inverse agonists AM251, rimonabant, and AM6538 suppressed basal signaling most and blocked CP55,940 most potently, whereas the neutral antagonists AM4113, AM6545, and AM6527 reduced constitutive activity significantly less, consistent with previously reported weaker or absent inverse-agonism (Cluny et al., 2010; Sink et al., 2009, 2008). AM4113 was the clear outlier, blocking Gαi1 only partially and leaving β-arrestin 2 recruitment largely intact, while AM6545 and AM6527 blocked both pathways.

Several of these efficacy assignments differ from prior classifications, perhaps reflecting the readout rather than the compound. MDMB-FUB, a full agonist at both receptors by fluorometric membrane-potential assay, and 2-AG, an established full agonist at CB2 by [³⁵S]GTPγS, both behaved as only partial agonists at CB2 Gαi1 here, while WIN55,212-2, classified as a nonselective full agonist, showed only partial efficacy at CB2. Because Gαi1 engagement BRET reports immediate receptor transducer coupling rather than an amplified second messenger, it is expected to return lower apparent efficacies than cAMP or [³⁵S]GTPγS readouts. In addition, normalizing every ligand to CP55,940 within the same assay places these compounds on a directly comparable scale.

There are some limitations to our conclusions. The measurements were made in a recombinant HEK293T system on two transducers, Gαi1 and β-arrestin 2, and extension to native tissues and to additional transducers was not assessed. Because CB1 exhibits constitutive activity in our assay, neutral antagonists can register a small reduction in basal signaling; separating a neutral antagonist from a weakly inverse agonist dose response would require additional controls or lower-expression systems. In addition, several β-arrestin 2 dose response curves did not reach a plateau, so their Emax and bias values represent lower bounds and are reported as not determined.

Nevertheless, the interpretations are supported by the uniform assay design, the normalization to CP55,940, and the comparison with established references, such as Δ⁹-THC. Future work with validation in native tissue, will be important to confirm and extend these findings.

## Conclusions

By profiling a broad cannabinoid panel against a CP55,940 reference under a uniform BRET assay approach, this study provides a directly comparable efficacy ranking of Gαi1 engagement and β-arrestin 2 recruitment at CB1 and CB2. Two patterns dominated: a widespread loss of agonist efficacy from CB1 to CB2, and pronounced β-arrestin 2 recruitment that was largely confined to CB1. Despite pronounced differences in β-arrestin 2 recruitment across the panel, most ligands showed no meaningful differences in their bias factors, suggesting that these efficacy differences did not translate into transducer bias in the absence of corresponding shifts in potency.

We were able to resolve antagonists by their degree of inverse agonism separating neutral antagonism from inverse agonism: inverse agonists suppressed constitutive CB1 activity more than neutral antagonists did. Several β-arrestin 2 dose response curves did not reach a plateau to provide EC50 values, thus yielding no conclusive corresponding Emax and bias factors. Although validation in native tissue will be needed to confirm and extend these *in vitro* pharmacological findings, overall, this uniform, CP55,940-normalized dataset offers a reference framework for understanding how cannabinoid structure governs receptor efficacy across the CB1 and CB2 signaling axes.

## Authorship Contribution

**Soo Jung Oh**: Data curation, Formal analysis, Investigation, Visualization, Writing, **Chrisotpher Lucaj**: Data curation, Formal analysis, Investigation, **Kiera Truong**: Data curation, Formal analysis, Investigation, **Ruiru Guo**: Data curation, Formal analysis, Investigation, **Hayato Umaoka**: Investigation, **Christos Iliopoulos-Tsoutsouvas**: Data curation, Formal analysis, Resources, **Maria Gerasi**: Resources, **Markos-Orestis Georgiadis**: Resources, **Lipin Ji**: Resources, **Spyros P Nikas**: Data curation, Formal analysis, Resources, **Alexandros Makriyannis**: Data curation, Formal analysis, Funding acquisition, Resources, Supervision, **Hideaki Yano**: Conceptualization, Data curation, Formal analysis, Funding acquisition, Resources, Supervision, Writing

## Acknowledgements

This work is supported by T32 training grant in SUD (S.J.O.), DA009158 (A.M.), and Northeastern University Startup Funds (H.Y.).

---

**Supplementary Table 1.**
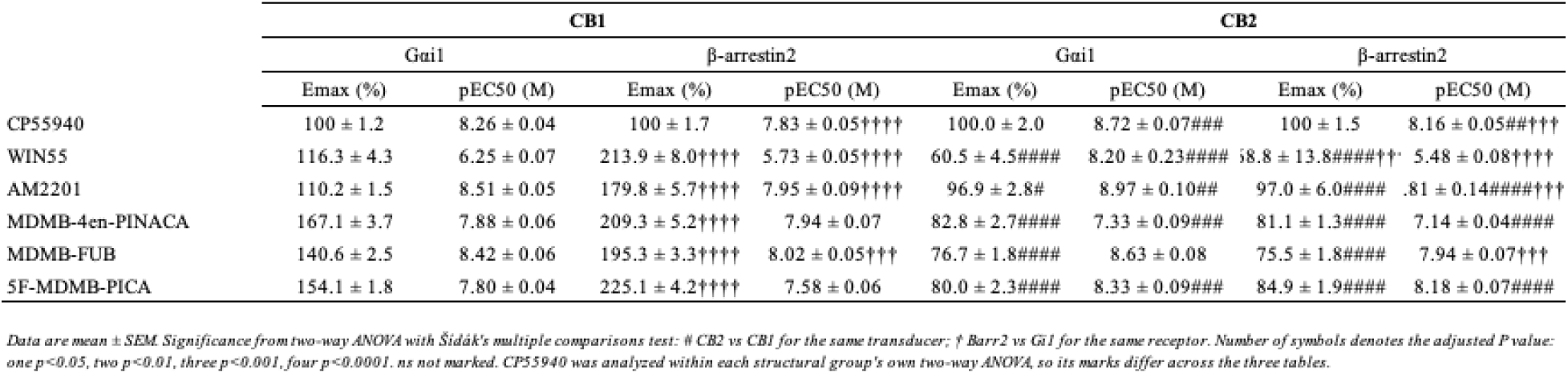
Comparison of SCRAs.

**Supplementary Table 2.**
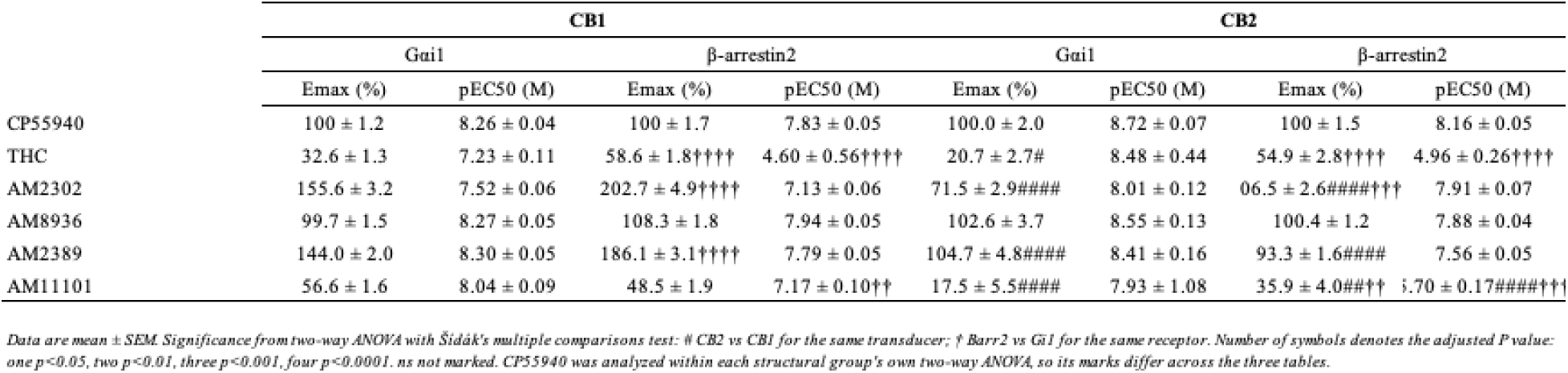
Comparison of THC Analogues.

**Supplementary Table 3.**
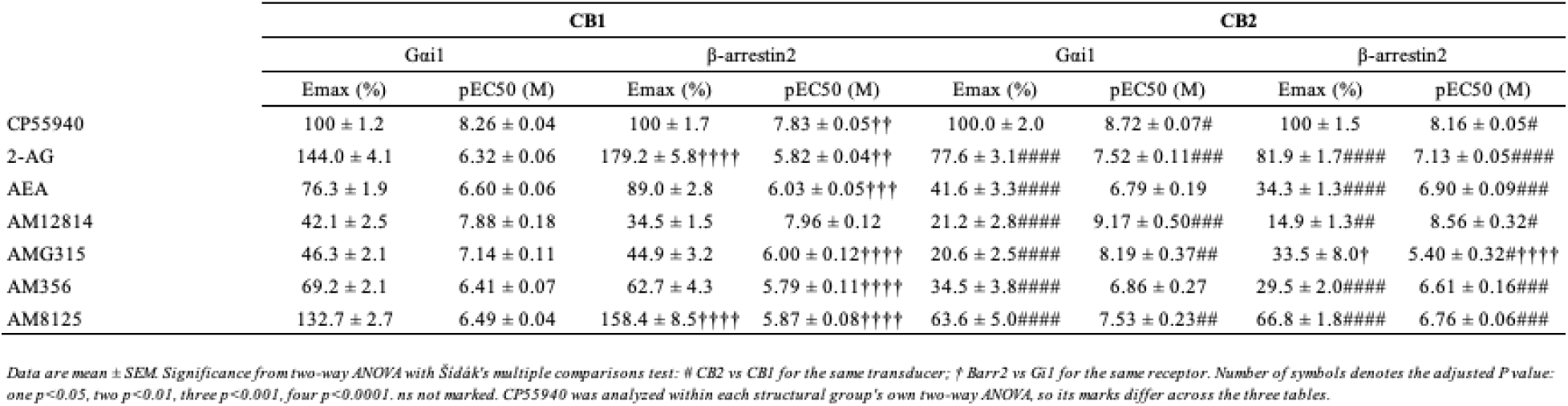
Comparison of Endocannabinoid Analogues.

## References

1. Abadji, V., Lin, S., Taha, G., Griffin, G., Stevenson, L.A., Pertwee, R.G., Makriyannis, A., 1994. (R)-methanandamide: a chiral novel anandamide possessing higher potency and metabolic stability. J. Med. Chem. 37, 1889–1893. 10.1021/jm00038a020

2. Atwood, B.K., Huffman, J., Straiker, A., Mackie, K., 2010. JWH018, a common constituent of “Spice” herbal blends, is a potent and efficacious cannabinoid CB receptor agonist. Br. J. Pharmacol. 160, 585–593. 10.1111/j.1476-5381.2009.00582.x

3. Banister, S.D., Longworth, M., Kevin, R., Sachdev, S., Santiago, M., Stuart, J., Mack, J.B.C., Glass, M., McGregor, I.S., Connor, M., Kassiou, M., 2016. Pharmacology of Valinate and tert-Leucinate Synthetic Cannabinoids 5F-AMBICA, 5F-AMB, 5F-ADB, AMB-FUBINACA, MDMB-FUBINACA, MDMB-CHMICA, and Their Analogues. ACS Chem. Neurosci. 7, 1241–1254. 10.1021/acschemneuro.6b00137

5. Banister, S.D., Stuart, J., Kevin, R.C., Edington, A., Longworth, M., Wilkinson, S.M., Beinat, C., Buchanan, A.S., Hibbs, D.E., Glass, M., Connor, M., McGregor, I.S., Kassiou, M., 2015. Effects of bioisosteric fluorine in synthetic cannabinoid designer drugs JWH-018, AM-2201, UR-144, XLR-11, PB-22, 5F-PB-22, APICA, and STS-135. ACS Chem. Neurosci. 6, 1445–1458. 10.1021/acschemneuro.5b00107

6. Caulkins, J.P., 2024. Changes in self-reported cannabis use in the United States from 1979 to 2022. Addiction 119, 1648–1652. 10.1111/add.16519

7. Chen, J., Li, F., Lee, J., Manirujjaman, M., Zhang, L., Song, Z.-H., McClain, C., Feng, W., 2024. Peripherally Restricted CB1 Receptor Inverse Agonist JD5037 Treatment Exacerbates Liver Injury in MDR2-Deficient Mice. Cells 13, 1101. 10.3390/cells13131101

8. Cinar, R., Iyer, M.R., Kunos, G., 2020. The therapeutic potential of second and third generation CB1R antagonists. Pharmacol. Ther. 208, 107477. 10.1016/j.pharmthera.2020.107477

9. Cluny, N., Vemuri, V., Chambers, A., Limebeer, C., Bedard, H., Wood, J., Lutz, B., Zimmer, A., Parker, L., Makriyannis, A., Sharkey, K., 2010. A novel peripherally restricted cannabinoid receptor antagonist, AM6545, reduces food intake and body weight, but does not cause malaise, in rodents. Br. J. Pharmacol. 161, 629–642. 10.1111/j.1476-5381.2010.00908.x

10. Di Marzo, V., De Petrocellis, L., 2012. Why do cannabinoid receptors have more than one endogenous ligand? Philos. Trans. R. Soc. B Biol. Sci. 367, 3216–3228. 10.1098/rstb.2011.0382

11. Felder, C.C., Joyce, K.E., Briley, E.M., Mansouri, J., Mackie, K., Blond, O., Lai, Y., Ma, A.L., Mitchell, R.L., 1995. Comparison of the pharmacology and signal transduction of the human cannabinoid CB1 and CB2 receptors. Mol. Pharmacol. 48, 443–450.

12. Gonsiorek, W., Lunn, C., Fan, X., Narula, S., Lundell, D., Hipkin, R.W., 2000. Endocannabinoid 2-arachidonyl glycerol is a full agonist through human type 2 cannabinoid receptor: antagonism by anandamide. Mol. Pharmacol. 57, 1045– 1050.

13. Gueye, A.B., Pryslawsky, Y., Trigo, J.M., Poulia, N., Delis, F., Antoniou, K., Loureiro, M., Laviolette, S.R., Vemuri, K., Makriyannis, A., Le Foll, B., 2016. The CB1 Neutral Antagonist AM4113 Retains the Therapeutic Efficacy of the Inverse Agonist Rimonabant for Nicotine Dependence and Weight Loss with Better Psychiatric Tolerability. Int. J. Neuropsychopharmacol. 19, pyw068. 10.1093/ijnp/pyw068

14. Hua, T., Vemuri, K., Pu, M., Qu, L., Han, G.W., Wu, Y., Zhao, S., Shui, W., Li, S., Korde, A., Laprairie, R.B., Stahl, E.L., Ho, J.-H., Zvonok, N., Zhou, H., Kufareva, I., Wu, B., Zhao, Q., Hanson, M.A., Bohn, L.M., Makriyannis, A., Stevens, R.C., Liu, Z.-J., 2016. Crystal Structure of the Human Cannabinoid Receptor CB1. Cell 167, 750–762.e14. 10.1016/j.cell.2016.10.004

15. Ibsen, M.S., Connor, M., Glass, M., 2017. Cannabinoid CB1 and CB2 Receptor Signaling and Bias. Cannabis Cannabinoid Res. 2, 48–60. 10.1089/can.2016.0037

16. Ibsen, M.S., Finlay, D.B., Patel, M., Javitch, J.A., Glass, M., Grimsey, N.L., 2019. Cannabinoid CB1 and CB2 Receptor-Mediated Arrestin Translocation: Species, Subtype, and Agonist-Dependence. Front. Pharmacol. 10. 10.3389/fphar.2019.00350

17. Idris, A.I., van ’t Hof, R.J., Greig, I.R., Ridge, S.A., Baker, D., Ross, R.A., Ralston, S.H., 2005. Regulation of bone mass, bone loss and osteoclast activity by cannabinoid receptors. Nat. Med. 11, 774–779. 10.1038/nm1255

18. Järbe, T.U.C., Tai, S., LeMay, B.J., Nikas, S.P., Shukla, V.G., Zvonok, A., Makriyannis, A., 2012. AM2389, a high-affinity, in vivo potent CB1-receptor-selective cannabinergic ligand as evidenced by drug discrimination in rats and hypothermia testing in mice. Psychopharmacology (Berl.) 220, 417–426. 10.1007/s00213-011-2491-1

19. Krotulski, A.J., Cannaert, A., Stove, C., Logan, B.K., 2021. The next generation of synthetic cannabinoids: Detection, activity, and potential toxicity of pent-4en and but-3en analogues including MDMB-4en-PINACA. Drug Test. Anal. 13, 427–438. 10.1002/dta.2935

20. Lan, R., Liu, Q., Fan, P., Lin, S., Fernando, S.R., McCallion, D., Pertwee, R., Makriyannis, A., 1999. Structure-activity relationships of pyrazole derivatives as cannabinoid receptor antagonists. J. Med. Chem. 42, 769–776. 10.1021/jm980363y

21. Landsman, R.S., Burkey, T.H., Consroe, P., Roeske, W.R., Yamamura, H.I., 1997. SR141716A is an inverse agonist at the human cannabinoid CB1 receptor. Eur. J. Pharmacol. 334, R1–2. 10.1016/s0014-2999(97)01160-6

22. Malamas, M.S., Raghav, J.G., Ma, X., Honrao, C., Wood, J.T., Benchama, O., Zhou, H., Mallipeddi, S., Makriyannis, A., 2018. Oximes short-acting CB1 receptor agonists. Bioorg. Med. Chem. 26, 4963–4970. 10.1016/j.bmc.2018.08.003

23. Malan, T.P., Ibrahim, M.M., Lai, J., Vanderah, T.W., Makriyannis, A., Porreca, F., 2003. CB2 cannabinoid receptor agonists: pain relief without psychoactive effects? Curr. Opin. Pharmacol. 3, 62–67. 10.1016/s1471-4892(02)00004-8

24. Munro, S., Thomas, K.L., Abu-Shaar, M., 1993. Molecular characterization of a peripheral receptor for cannabinoids. Nature 365, 61–65. 10.1038/365061a0

25. Namkung, Y., Le Gouill, C., Lukashova, V., Kobayashi, H., Hogue, M., Khoury, E., Song, M., Bouvier, M., Laporte, S.A., 2016. Monitoring G protein-coupled receptor and β-arrestin trafficking in live cells using enhanced bystander BRET. Nat. Commun. 7, 12178. 10.1038/ncomms12178

26. Nikas, S.P., Alapafuja, S.O., Papanastasiou, I., Paronis, C.A., Shukla, V.G., Papahatjis, D.P., Bowman, A.L., Halikhedkar, A., Han, X., Makriyannis, A., 2010. Novel 1′,1′-Chain Substituted Hexahydrocannabinols: 9β-Hydroxy-3-(1-hexyl-cyclobut-1-yl)-hexahydrocannabinol (AM2389) a Highly Potent Cannabinoid Receptor 1 (CB1) Agonist. J. Med. Chem. 53, 6996–7010. 10.1021/jm100641g

27. Olsen, R.H.J., DiBerto, J.F., English, J.G., Glaudin, A.M., Krumm, B.E., Slocum, S.T., Che, T., Gavin, A.C., McCorvy, J.D., Roth, B.L., Strachan, R.T., 2020. TRUPATH, an open-source biosensor platform for interrogating the GPCR transducerome. Nat. Chem. Biol. 16, 841–849. 10.1038/s41589-020-0535-8

28. Papanastasiou, I.P., Georgiadis, M.-O., Iliopoulos-Tsoutsouvas, C., Paronis, C.A., Brust, C.A., Tran, N.K., Ji, L., Ma, X., Wood, J.T., Zvonok, N., Tong, F., Bohn, L. M., Nikas, S.P., Makriyannis, A., 2022. Improved cyclobutyl nabilone analogs as potent CB1 receptor agonists. Eur. J. Med. Chem. 230, 114027. 10.1016/j.ejmech.2021.114027

30. Pertwee, R.G., 2008. The diverse CB1 and CB2 receptor pharmacology of three plant cannabinoids: delta9-tetrahydrocannabinol, cannabidiol and delta9-tetrahydrocannabivarin. Br. J. Pharmacol. 153, 199–215. 10.1038/sj.bjp.0707442

31. Rinaldi-Carmona, M., Barth, F., Héaulme, M., Shire, D., Calandra, B., Congy, C., Martinez, S., Maruani, J., Néliat, G., Caput, D., 1994. SR141716A, a potent and selective antagonist of the brain cannabinoid receptor. FEBS Lett. 350, 240–244. 10.1016/0014-5793(94)00773-x

32. Sink, K.S., McLaughlin, P.J., Wood, J.A.T., Brown, C., Fan, P., Vemuri, V.K., Peng, Y., Olszewska, T., Thakur, G.A., Makriyannis, A., Parker, L.A., Salamone, J.D., 2008. The novel cannabinoid CB1 receptor neutral antagonist AM4113 suppresses food intake and food-reinforced behavior but does not induce signs of nausea in rats. Neuropsychopharmacol. Off. Publ. Am. Coll. Neuropsychopharmacol. 33, 946– 955. 10.1038/sj.npp.1301476

33. Sink, K.S., Vemuri, V.K., Wood, J., Makriyannis, A., Salamone, J.D., 2009. Oral bioavailability of the novel cannabinoid CB1 antagonist AM6527: effects on food-reinforced behavior and comparisons with AM4113. Pharmacol. Biochem. Behav. 91, 303–306. 10.1016/j.pbb.2008.07.013

34. Tsang, C.C., Giudice, M.G., 2016. Nabilone for the Management of Pain. Pharmacotherapy 36, 273–286. 10.1002/phar.1709

35. Turcotte, C., Blanchet, M.-R., Laviolette, M., Flamand, N., 2016. The CB2 receptor and its role as a regulator of inflammation. Cell. Mol. Life Sci. CMLS 73, 4449–4470. 10.1007/s00018-016-2300-4

36. Wang, T., Li, Z., Cvijic, M.E., Zhang, L., Sum, C.S., 2004. Measurement of cAMP for Gαs- and Gαi Protein-Coupled Receptors (GPCRs), in: Markossian, S., Grossman, A., Baskir, H., Arkin, M., Auld, D., Austin, C., Baell, J., Brimacombe, K., Chung, T.D.Y., Coussens, N.P., Dahlin, J.L., Devanarayan, V., Foley, T.L., Glicksman, M., Gorshkov, K., Grotegut, S., Hall, M.D., Hoare, S., Inglese, J., Iversen, P.W., Lal-Nag, M., Li, Z., Manro, J.R., McGee, J., Norvil, A., Pearson, M., Riss, T., Saradjian, P., Sittampalam, G.S., Tarselli, M.A., Trask, O.J., Weidner, J.R., Wildey, M.J., Wilson, K., Xia, M., Xu, X. (Eds.), Assay Guidance Manual. Eli Lilly & Company and the National Center for Advancing Translational Sciences, Bethesda (MD).

37. World Drug Report, 2026. World Drug Report 2026 [WWW Document]. U. N. Off. Drugs Crime. URL //www.unodc.org/unodc/en/data-and-analysis/world-drug-report-2026-highlights.html (accessed 9.16.26).

38. Yang, H., Park, M., Lee, J.H., Kim, B., Moon, C.S., Bae, S., Kim, Y., Lee, H.-J., Park, C.-Y., 2024. New peripherally-restricted CB1 receptor antagonists, PMG-505-010 and -013 ameliorate obesity-associated NAFLD and fibrosis. Biomed. Pharmacother. Biomedecine Pharmacother. 180, 117501. 10.1016/j.biopha.2024.117501

39. Zou, S., Kumar, U., 2018. Cannabinoid Receptors and the Endocannabinoid System: Signaling and Function in the Central Nervous System. Int. J. Mol. Sci. 19, 833. 10.3390/ijms19030833

